# Neoadjuvant chemotherapy induces transient, regimen-specific immune–stromal reprogramming in pancreatic ductal adenocarcinoma

**DOI:** 10.64898/2026.09.07.749769

**Authors:** Elahe Minaei, Dezerae Cox, Amarinder Singh Thind, Gary Tincknell, Leo A. Coleman, Samantha J. Wade, Naila Islam, Dale Waring, Mona Ahuja, Joanne La Malfa, Koroush S Haghighi, Marie Ranson, Morteza Aghmesheh, Ronald Sluyter, Eileen M. O’Reilly, Kara L. Vine-Perrow

## Abstract

**Background:** Neoadjuvant chemotherapy (NAC) is increasingly incorporated into the management of pancreatic ductal adenocarcinoma (PDAC), yet how treatment regimen and timing reshape the tumour-immune microenvironment remains poorly defined.

**Methods:** Targeted immune transcriptomic profiling was performed using NanoString on spatially annotated tumour, stromal, and immune-enriched regions from resected PDAC specimens from 23 patients, including NAC-treated and treatment-naïve cohorts. Key findings were validated and spatially localised using Xenium *in situ* transcriptomics.

**Results:** NAC induced extensive transcriptional remodelling of the tumour microenvironment in a regimen- and time-dependent manner. Tumours resected shortly after FOLFIRINOX exhibited coordinated upregulation and strong spatial coupling of the *CXCL12–CXCR4* axis. These correlations were attenuated or reversed at longer post-treatment intervals and following gemcitabine-based therapy. Disrupted stromal coordination of the *CCL2–CCR2* axis further highlighted context-dependent effects of chemotherapy on chemokine signalling. Across regions, *NT5E* (CD73) expression inversely correlated with *CD8A*, consistent with spatially restricted T cell exclusion.

**Conclusions:** NAC dynamically reprograms the immune–stromal landscape of PDAC in a regimen- and timing-dependent manner, revealing a transient post-chemotherapy window of immune remodelling. The clinical significance of this phenomenon requires further investigation to determine whether temporally optimised integration of immunomodulatory and stroma-targeted therapies with NAC can improve patient outcomes.

**Graphical abstract:** 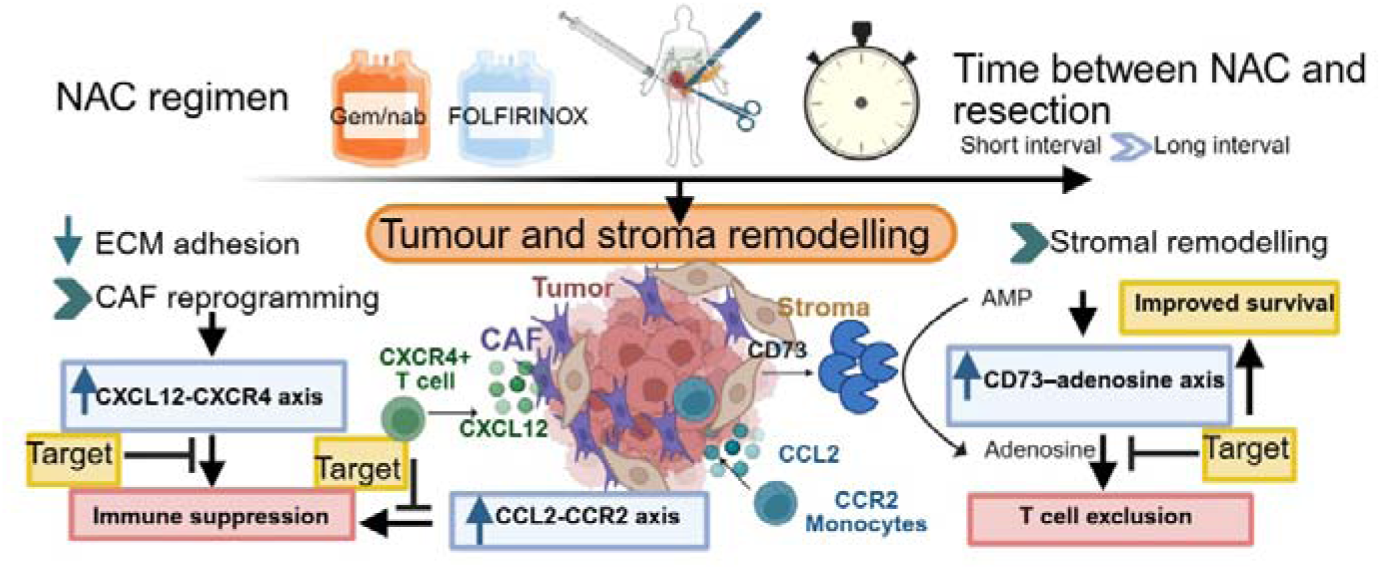

## Background

Pancreatic ductal adenocarcinoma (PDAC) remains one of the deadliest cancers, with a five-year survival rate of less than 12%1, largely due to late diagnosis, rapid metastatic progression and limited responsiveness to existing systemic therapies (1). Neoadjuvant chemotherapy (NAC), including FOLFIRINOX (FOLinic acid, 5-Flurouracil, IRINotecan, OXaliplatin) and gemcitabine-based regimens, is increasingly incorporated into the management of borderline resectable and locally advanced disease to improve resectability and early disease control (2). Despite modest improvements in clinical outcomes, long-term survival remains poor with such treatments (3). PDAC is characterised by a dense, fibrotic and immune-excluding tumour microenvironment (TME), which contributes to profound resistance to immunotherapy; however, how NAC reshapes this immune–stromal landscape remains incompletely understood. Increasing attention has therefore focused on the capacity of conventional cytotoxic therapies to remodel the tumour immune microenvironment and influence responsiveness to immunomodulatory interventions with the aim of improving patient outcomes (4).

Although chemotherapy has historically been regarded as cytotoxic and immunosuppressive (4), accumulating evidence indicates that its immunological effects are highly context dependent. Under specific conditions, certain agents can stimulate anti-tumour immunity by either inducing immunogenic cell death or immune–stromal interactions within the TME (5). Oxaliplatin and anthracyclines for example, can trigger immunogenic cell death via the release of damage-associated molecular patterns (DAMPs), including calreticulin, adenosine 5’-triphosphate and high mobility group box 1, thereby promoting dendritic cell activation and cytotoxic T cell priming (6, 7). In contrast, other chemotherapies may reinforce immune suppression or immune evasion (8). Gemcitabine, while not a classical inducer of immunogenic cell death, has been shown to enhance CD8⁺ T cell infiltration and increase PD-L1 expression, potentially converting immunologically “cold” tumours into more inflamed states that may be permissive to immune checkpoint blockade (9).

Despite these observations, the immunological consequence of chemotherapy in PDAC remain inconsistent across studies and appear to depend on treatment composition, timing of tissue sampling, concomitant radiotherapy and patient-specific factors. For example, multiplex GeoMx protein analysis and NanoString tumour RNA profiling demonstrated that neoadjuvant FOLFIRINOX alone increased PD1 and granzyme B expression and enriched interferon-related pathways, consistent with favourable immune remodelling, whereas the addition of stereotactic body radiotherapy reversed these effects and was associated with depletion of CD3⁺ and CD4⁺ T cells and downregulation of immune checkpoint molecules (10). Similarly, single-cell RNA sequencing studies have reported reduced immune checkpoint expression and diminished T cell–tumour interactions following chemotherapy, suggesting persistent immune evasion despite treatment (11). In contrast single-nucleus RNA sequencing analyses have shown enrichment of CD8⁺ T cells, neurotropic cancer-associated fibroblasts (CAFs) and mesenchymal tumour populations following neoadjuvant FOLFIRINOX, highlighting the concurrent induction of immune activation and stromal reprogramming (12). More recently, sex-specific differences in immune remodelling following gemcitabine-based chemoradiotherapy were reported in PDAC patients from the PREOPANC trial (13), underscoring the influence of host factors on therapy-induced TME responses (13).

Collectively, these studies underscore the complexity of chemotherapy-induced immune remodelling in PDAC but also reveal key gaps in our current understanding. In particular, how neoadjuvant chemotherapy reshapes immune–stromal signalling in a spatially resolved manner, and whether these effects vary according to a specific treatment regimen and timing relative to surgery, remains largely unexplored. To address this, we employed spatially annotated NanoString immune profiling in combination with Xenium *in situ* gene expression analysis to characterise immune and stromal remodelling in treatment-naïve (TN) and neoadjuvant chemotherapy (NAC)–treated PDAC specimens. By comparing tumour-rich, stromal-rich, immune-rich and tumour-adjacent normal tissue compartments, we aimed to define spatial and molecular features of the TME associated with neoadjuvant therapy. These insights provide a framework for understanding how chemotherapy-induced immune–stromal reprogramming may inform the rational timing and design of combination therapeutic strategies in PDAC.

## Methods

### Study population and sample collection

This study was approved by the University of Wollongong and Illawarra Shoalhaven Local Health District Human Research Ethics Committee (UOW/ISLHD HREC 2020/ETH03297) and conducted in accordance with the Declaration of Helsinki and the National Statement on Ethical Conduct in Human Research (NHMRC, Australia). Slides and formalin-fixed, paraffin embedded (FFPE) PDAC blocks from 23 patients who received neoadjuvant chemotherapy (NAC = 11) or were treatment-naïve (TN = 12) were retrieved from Health Precincts Biobank at Prince of Wales Hospital (Sydney, Australia) and NSW Health Pathology at Wollongong Hospital (Wollongong, Australia). All specimens were de-identified prior to analysis. As the study involved retrospective archival biospecimens, the Human Research Ethics Committee approved a waiver of consent. Hematoxylin and eosin (H&E) stained slides were examined by pathologists under a light microscope to identify and annotate specific target areas. Tumour-rich regions were defined as areas with a high percentage of malignant cells, ideally 100%, though regions with 80–100% cancer cells were acceptable. These regions excluded areas of necrosis, inflammation, fibrosis and normal ductal tissue. Stroma-rich regions were selected based on a predominance of stromal tissue, such as fibroblasts and collagen, with less than 15% malignant cells and minimal immune cell infiltration. Immune cell-rich areas were identified as peri-tumoural and inter-tumoural zones with a high density of immune cells and low cancer cell content. Normal parenchymal tissue was also annotated for comparative analysis. The marked areas on the H&E slides were matched to corresponding paraffin blocks and three or four tissue cores (1 mm diameter, ∼4 mm depth) were extracted from each targeted area.

### RNA extraction and Nanostring gene expression assay

Tissue cores were deparaffinised and homogenised to facilitate RNA extraction from FFPE specimens. Tumour nucleic acids were extracted using the AllPrep DNA/RNA FFPE Kit (cat# 80234, Qiagen, Hilden, Germany), following the manufacturer’s protocol to ensure optimal yield and integrity. The deparaffinisation process involved xylene wash, followed by rehydration through a graded ethanol series, which allows for an efficient removal of paraffin. Homogenisation of the tissue was carried out using a rotor-stator homogeniser (Omni tissue homogeniser, Revvity, GA, USA) to ensure uniform disruption of the sample, to maximise nucleic acid recovery.

RNA extraction involved proteinase K digestion for lysis, followed by binding nucleic acids to a silica membrane for isolation of high-purity RNA. Purified RNA was quantified and subjected to quality control (QC) assessment. RNA integrity and purity were determined using the Invitrogen Qubit RNA IQ Assay (Thermo Fisher Scientific, Waltham, USA), with acceptable RNA integrity (IQ) values set between 5 and 10 and A260/280 nm absorbance ratios ranging from 1.8 to 2 measured by Nanodrop 2000c Spectrometer (Thermo Fisher Scientific). Only RNA samples meeting these QC thresholds were included in subsequent gene expression analyses to ensure reliable downstream data. NanoString PanCancer Immune Profiling (740 genes) was performed and analysed after standard QC, normalisation and batch correction; differential expression used DESeq2 with false discovery rate (method in Supplementary materials).

### Gene set enrichment using over-representation analysis

DEGs were identified from RNA-seq data, with significance thresholds set based on p-value < 0.01 (given the exploratory nature of this study and the limited sample size, an FDR threshold of 0.2-0.3 (may vary per analysis) was applied, consistent with practices in underpowered discovery analyses (14)) and log₂ fold change > 1. The resulting list of significant genes was used as an input for gene set enrichment analysis using Enricher, a tool that performs over-representation analysis across a wide variety of gene set libraries. The background was set based on the total number of genes in the NanoString panel. Enrichment was analysed across several relevant libraries, including KEGG (2021 Human), MSigDB Hallmark and BioPlanet (2019) databases. The databases calculate enrichment using a combination of the Fisher exact test, a Z-score and a combined score, which is computed by multiplying the log-transformed p-value by the Z-score of the deviation from the expected rank. Results were filtered based on adjusted p-value < 0.05 for significance. Top enriched terms were visualised using in-house R script.

### Xenium *in situ* gene expression assay

Xenium *in situ* spatial transcriptomic profiling was performed as a fee-for-service by the Garvan Institute of Medical Research, (Sydney, Australia). Formalin-fixed paraffin-embedded (FFPE) tissue blocks were sectioned at 5 µm thickness and mounted onto Xenium slides. Spatial gene expression analysis was conducted using the Xenium Human Immuno-Oncology Profiling Panel targeting 380 genes. All subsequent experimental procedures, including tissue processing, probe hybridisation, amplification, imaging, and primary data generation, were performed by the service provider according to manufacturer-validated protocols using the Xenium Analyser (10x Genomics). Automated image acquisition and transcript detection were carried out on the Xenium Analyser. Image processing, transcript decoding, quality filtering, and cell segmentation were performed using the proprietary Xenium analysis pipeline. Quality-filtered spatial transcriptomic outputs, including transcript coordinates, feature-cell matrices, and cell boundary annotations, were generated by the service provider. Downstream spatial visualisation and differential gene expression analyses were conducted using Xenium Explorer software and Python (v3.9).

### Statistical analyses

Clinical and demographic variables between treatment-naïve and neoadjuvant chemotherapy groups were compared using Fisher’s Exact Test for categorical variables and the Mann–Whitney U test for continuous variables, with significance defined as p-value < 0.05. Over-representation analysis was performed using Enrichr across MSigDB Hallmark, KEGG and BioPlanet gene set libraries. Enrichment significance was calculated using Fisher’s exact test and results were ranked by adjusted p-values (< 0.05) and combined scores. For spatial transcriptomics data generated by Xenium (10× Genomics), region-specific expression correlations (e.g. ligand-receptor and immunosuppressive axes) were evaluated using Pearson’s correlation on spatially binned data (500 × 500 pixels per bin, ∼918 cells/bin). Statistical significance was determined using two-tailed p-values with 95% confidence intervals. Spatial interactions between cell types were assessed via neighbourhood enrichment analysis comparing observed co-localisation frequencies to randomised spatial distributions. All statistical analyses were conducted in R (v4.2.0) and Python (v3.9) using standard packages and custom scripts available from Zenodo (Geneva, Switzerland) (10.5281/zenodo.17157651). A p-value < 0.05 was considered statistically significant unless otherwise specified.

## Results

### Patient cohort and data collection

Patient clinical and demographic data are summarised in Table 1. A total of 39 samples from 23 patients were included in the immune profiling analysis following RNA quality assessment (IQ > 4) using Qubit and NanoString quality control, with batch and instrument effects removed. Of the 23 patients for whom tumour-, stroma-, and/or immune-rich region specimens were included, eight received FOLFIRINOX (between 2 and 8 cycles of neo-adjuvant chemotherapy) and three received gemcitabine/nab-paclitaxel (between 5-12 cycles). One patient who received FOLFIRINOX subsequently received concurrent chemoradiotherapy with capecitabine prior to surgery. The remaining 12 patients were treatment-naïve (TN). No significant differences were observed in age or gender distribution between NAC and TN groups. The interval between the last cycle of chemotherapy and surgery ranged from 2 to 13 weeks. One patient who received additional chemoradiotherapy underwent surgery 26 weeks after completion of neoadjuvant chemotherapy (patient clinical history is provided in the Supplementary Data file 1).

**Table 1.** Clinical and demographic PDAC patient data.

|  |  | **TN** | **NAC** | **P value** |
| --- | --- | --- | --- | --- |
| **Age at diagnosis**<br>**(Median, IQR)\*** |  | 67.5 (63, 70.8) | 67.0 (62.5, 72.5) | 1.00 |
| **Sex (%)** | Female | 5 | 8 | 0.28 |
|  | Male | 7 | 3 |  |
| **Tissues included**<br>**in the analysis** | Tumour (A) | 7 | 6 |  |
|  | Stroma (B) | 7 | 7 |  |
|  | Immune (C) | 7 | 5 |  |
|  | Normal (D) | 5 | 6 |  |
| **Pathological**<br>**TNM staging**<br>**(%)** | I | 3 (25) | 4 (36.4) | 0.58 |
|  | II | 6 (50) | 6 (54.5) |  |
|  | III | 3 (25) | 1 (9.1) |  |
| **Time from end**<br>**of neoadjuvant**<br>**therapy to**<br>**surgery (weeks)** | ≤ 4 | NA | 2 (18.2) |  |
|  | 4-12 | NA | 3 (27.3) |  |
|  | ≥12≤ | NA | 3 (27.3) |  |
|  | Unknown |  | 3 (27.3) |  |
| **Recurrence**<br>**(months)** | No | 4 (33.3) | 3 (37.3) | 0.30 |
|  | Yes | 8 (66.7) | 6 (54.5) |  |
|  | Unknown | 0 (0) | 2 (18.2) |  |

\*IQR, Interquartile Range: measure of statistical spread.

### Neoadjuvant chemotherapy induces regimen- and timing-dependent immune and stromal transcriptional remodelling in PDAC

Differential gene expression analysis across 39 pooled tissue samples from TN and NAC-treated patients was performed. Differential gene expression data are presented in Supplementary Data File 1, ordered from the lowest to the highest adjusted p-value (padj). Unsupervised hierarchical clustering of selected tumour-, immune- and stromal-associated genes revealed distinct separation between TN and NAC groups (Figure 1A), indicating widespread transcriptional reprogramming, following cytotoxic therapy. Comparative analysis revealed significant upregulation of chemotactic and immune-associated genes in NAC-treated tumours, including *CCL2, CXCL12, CD163, S100A12, CCL17*, and *THBS1* (thrombospondin 1) (Figure 1B). In contrast, several genes associated with extracellular matrix interactions, cell adhesion, survival, inflammation, and metabolic regulation including *ITGB4* (α6β4 integrin), *ITGA2* (CD49b), *PTGS2* (prostaglandin-endoperoxide synthase 2), BIRC5 (baculoviral IAP repeat containing 5), *PPARG* (peroxisome proliferator-activated receptor γ), and *NT5E* (CD73) were downregulated following NAC (Figure 1B). Expression of *CD8A*, a marker of cytotoxic T cells, was also increased in NAC-treated tumours.

**Figure 1.**
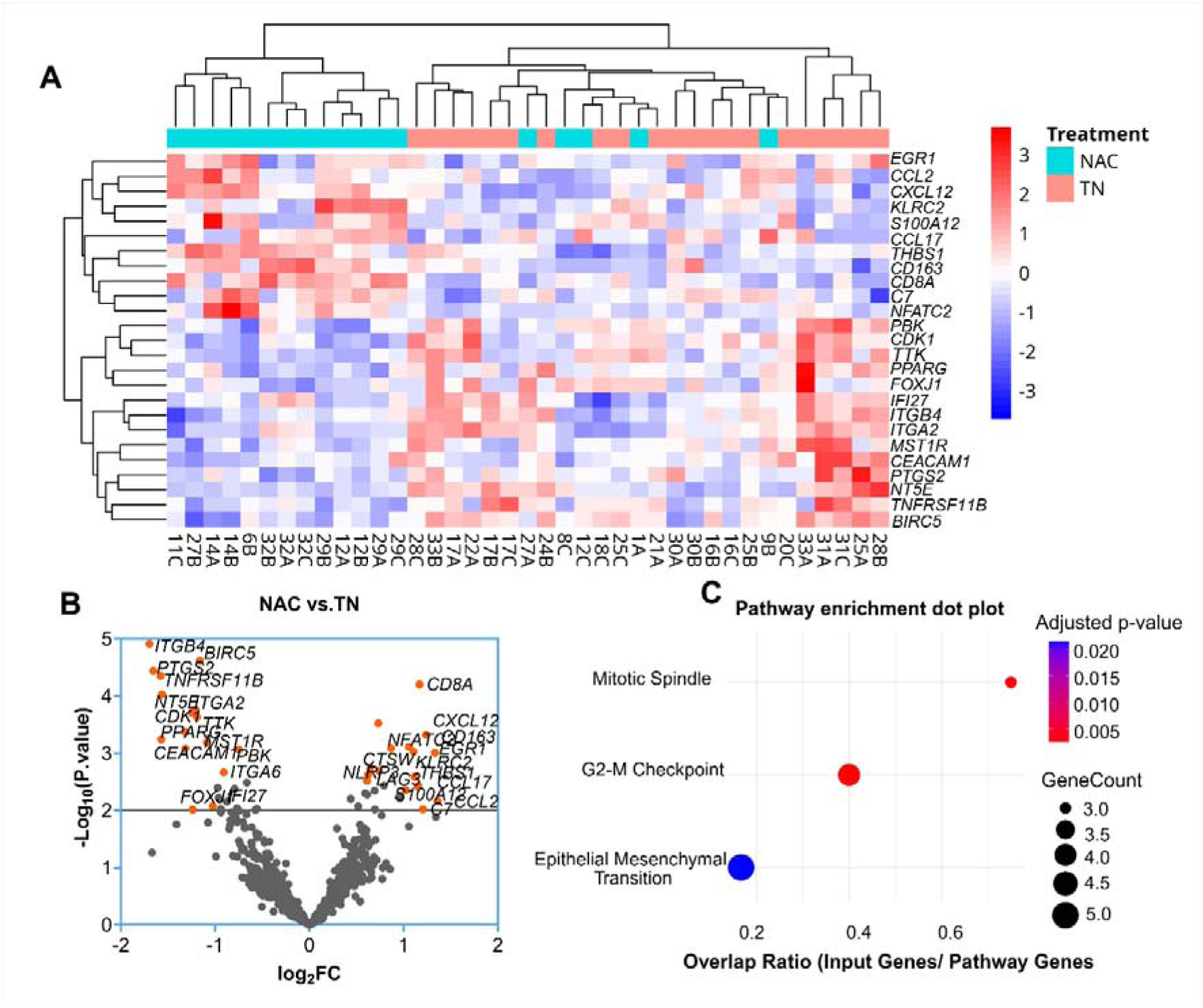
Pooled tissue NanoString gene expression and pathway analysis. **(A)** Unsupervised hierarchical clustering heatmap of top differentially expressed immune-related genes from NanoString analysis, comparing neoadjuvant chemotherapy (NAC)-treated (cyan) and treatment-naïve (TN, pink) PDAC patient samples. Rows represent genes and columns represent individual tissue samples. **(B)** Volcano plot showing log₂ fold change (x-axis) versus log₁₀ adjusted p-value (y-axis) for NAC vs. TN comparison. Key upregulated and downregulated genes are highlighted. **(C)** MSigDB Hallmark 2020 Pathway enrichment analysis of differentially expressed genes between NAC and TN groups. Dot size indicates the number of genes in each enriched pathway and dot colour reflects the adjusted p-value < 0.05.

Pathway enrichment analysis of differentially expressed genes using MSigDB Hallmark gene sets identified significant enrichment of Mitotic Spindle and G2–M Checkpoint pathways (adjusted p-value < 0.01), as well as enrichment of the Epithelial–Mesenchymal Transition pathway (adjusted p-value = 0.022) (Figure 1C). Reactome pathway analysis highlighted enrichment of Extracellular Matrix (ECM)–Related pathways, including non-integrin membrane ECM interactions, syndecan interactions, and laminin interactions (Supplementary Table S1). BioPlanet analysis revealed enrichment of transcriptional programs related to BDNF signalling, AP-1, and NFAT networks (Supplementary Table S2). Tissue-specific gene expression and pathway analyses across tumour, stromal, and immune-rich regions are presented in the Supplementary Material, Supplementary result and discussion section (This section includes Supplementary Figure S1 and Tables S3-S10).

Given the known biological and treatment heterogeneity of PDAC, spatial analyses were prioritised to identify conserved patterns of immune–stromal coordination rather than uniform expression changes across all cases.

### Spatial transcriptomics identifies stromal and myeloid compartments as key drivers of chemotherapy-associated immune suppression

#### Consistent cell density and high-quality spatial transcriptomic coverage across PDAC samples

Single-cell imaging using Xenium *in situ* gene expression was performed to characterise immune and stromal organisation in representative NAC-treated and treatment-naïve (TN) PDAC samples (Supplementary Table S11). Xenium data quality was consistent across samples, with comparable median cell densities and no evidence of transcript dropout, enabling downstream cell-type and spatial analyses (Figure 2A–C). Across all samples, the distribution of total transcripts per cell was right-skewed, with most cells expressing fewer than 100 transcripts. The mean transcript count per cell was 55.3 ± 49.8 (Figure 2A), while the mean number of unique transcripts per cell was 19.6 ± 18.7 (Figure 2B), indicating consistent transcript capture across samples and sufficient resolution for cell-type classification. Violin plots of cell density (cells per 100 µm²) across individual patient samples demonstrated broadly consistent and well-distributed cellular coverage across both NAC and TN tissues (Figure 2C). Unsupervised clustering using the Leiden algorithm on UMAP-reduced transcriptomic space identified 21 distinct cell clusters (Figure 2D). Cluster identities were assigned based on canonical marker gene expression visualised in a dot plot (Figure 2E), following automated annotation and subsequent manual refinement.

**Figure 2.**
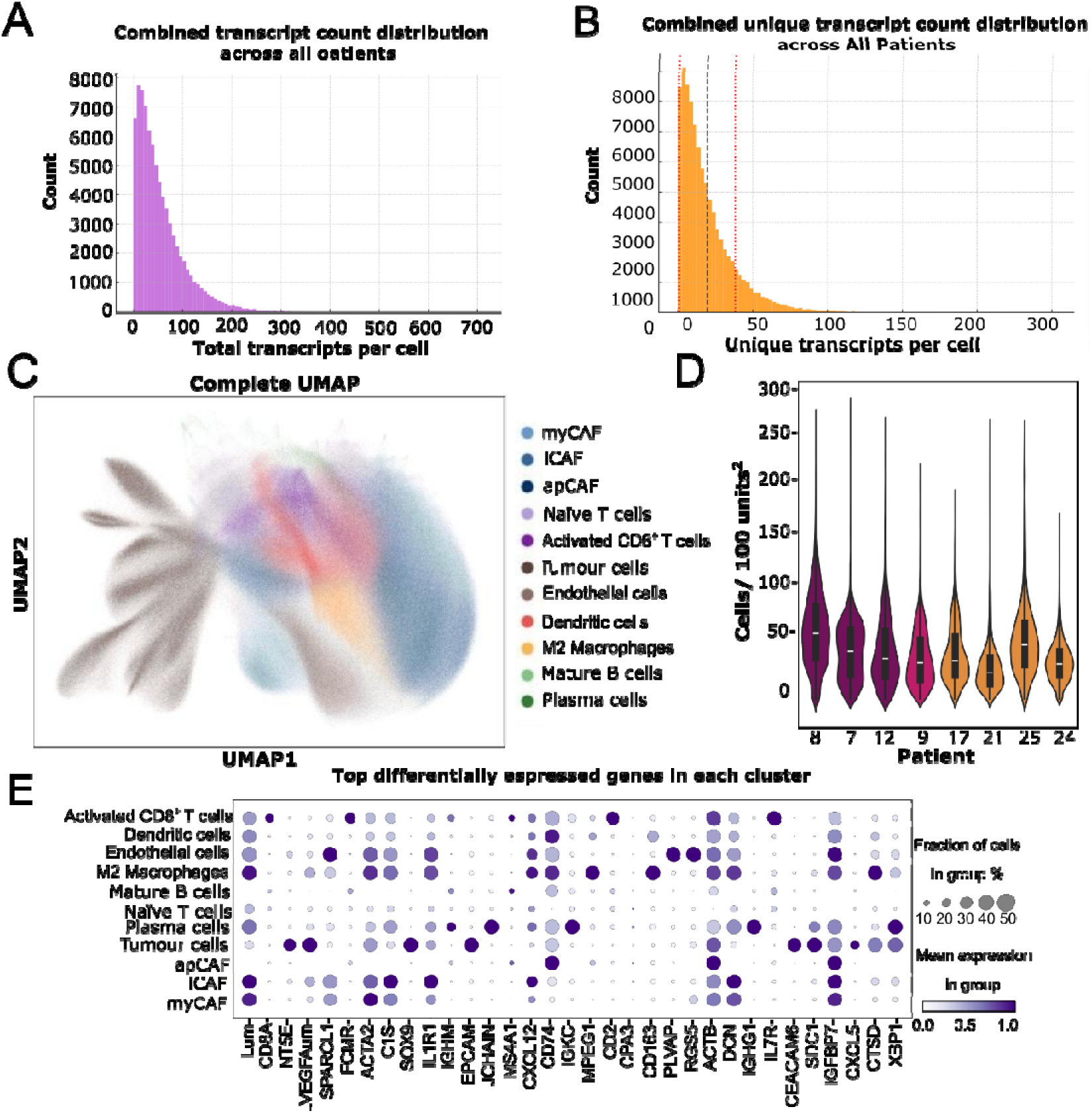
Quality control metrics and cell type annotation for Xenium spatial transcriptomic data from PDAC samples. (A) Distribution of gene counts per cell across all samples. (B) Histogram of unique transcript counts per cell across all patients. Dotted lines indicate QC thresholds for minimum (10 transcripts; black) and maximum (300 transcripts; red) filtering. (C) UMAP visualisation of all annotated cells coloured by cell type identity. (D) Violin plots of transcript count per cell stratified by sample and cell type. NAC-treated samples are shown in orange and TN samples in purple. (E) Dot plot of canonical marker gene expression used to guide manual cell type annotation, where dot size represents percent expressing cells and colour intensity reflects average expression.

#### Chemotherapy timing influences immune cell localisation and stromal association within the PDAC microenvironment

Spatial mapping of annotated cell types overlaid with pathology-defined regions of interest revealed treatment-associated differences in immune distribution and immune–stromal architecture (Figure 3A). Immune-rich regions were characterised by high densities of CD8⁺ T cells and dendritic cells, while stroma-rich regions were dominated by cancer-associated fibroblast populations. Tumour-rich regions displayed variable immune infiltration depending on treatment status and timing.

**Figure 3.**
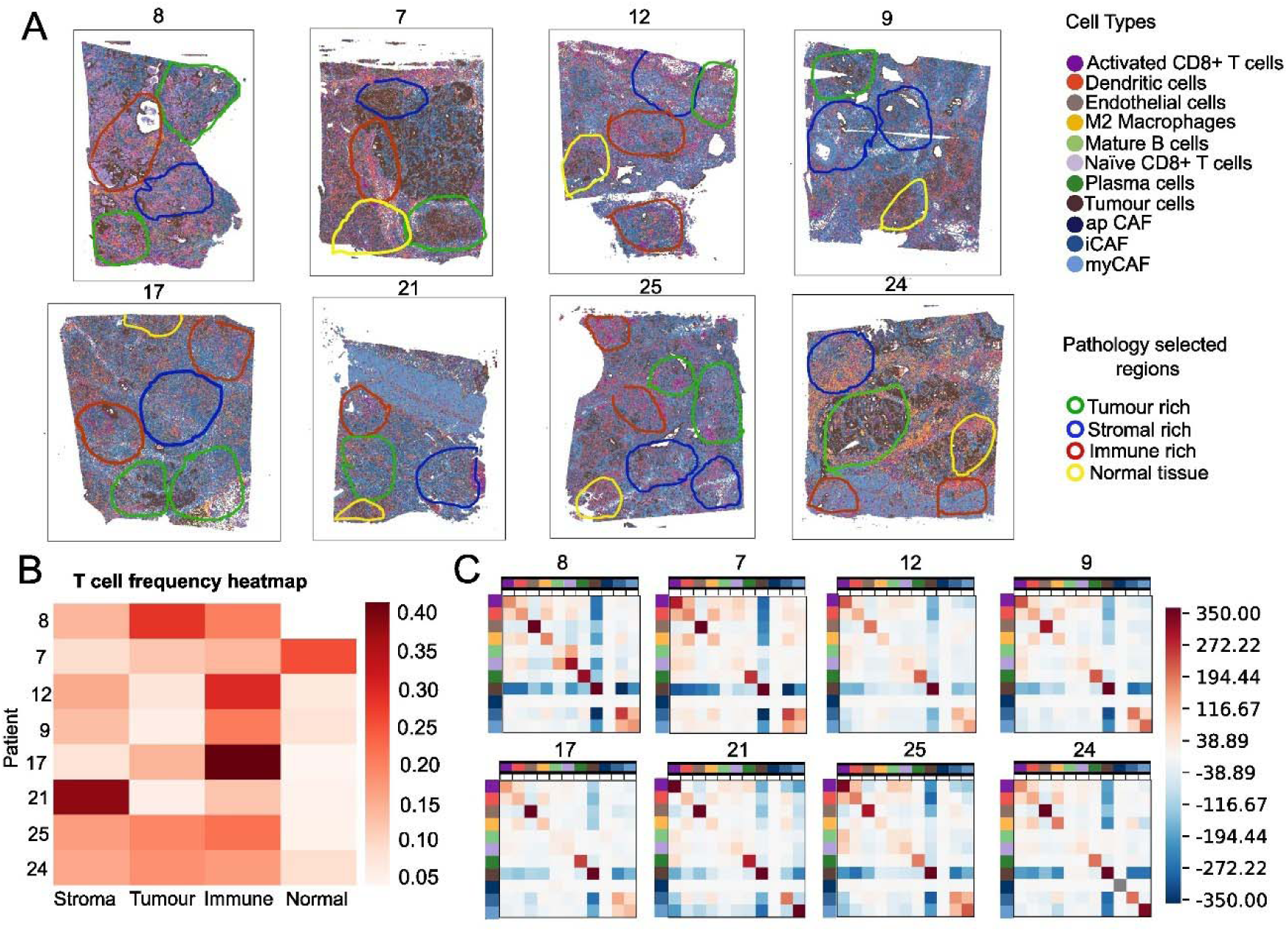
Spatial distribution, abundance and neighbourhood analysis of immune and stromal cells across pathology-defined PDAC tissue regions. (A) Xenium spatial maps from four NAC patients (8, 7, 12, 9) and four TN patients (17, 21, 25, 24). PDAC patients showing single-cell resolution cell type annotation overlaid with pathologist-defined tissue compartments: Tumour-rich (green), stroma-rich (blue), immune-rich (red) and adjacent normal tissue (yellow). Cell types were defined based on marker gene expression and include various immune, stromal and tumour cell populations as indicated in the legend. (B) Heatmap of T cell frequency (activated and naïve CD8⁺ T cells combined) across annotated tissue compartments for each patient. Frequencies represent the proportion of total cells in each compartment that are T cells. (C) Cell–cell neighbourhood interaction matrices computed for each patient. Colours represent the degree of spatial enrichment (red) or depletion (blue) of one cell type in the proximity of another, compared to a random background. The colour code on the top and left of the spatial enrichment graphs represents the cell types.

In NAC-treated patients 9 and 12 (both with 12-week intervals between chemotherapy and surgery), tumour-rich regions remained largely devoid of T cells. In contrast, patient 8, who underwent surgery three weeks after completing FOLFIRINOX, exhibited a more diffuse distribution of T cells across tumour, stromal, and immune-rich compartments.

Quantification of T cell frequency across tissue compartments showed that immune-rich regions generally harboured the highest T cell densities (Figure 3B), except for NAC patients 8 and 7 and TN patient 21. In patient 7, T cells were most abundant in adjacent normal tissue. In patient 8, the highest proportion of T cells was detected in tumour-rich regions.

#### Spatial coordination of immunosuppressive chemokine–receptor axes following neoadjuvant chemotherapy

Building on the NanoString findings, spatial transcriptomic analysis was performed to contextualise and localise the differential expression of key immune genes between NAC-treated and TN PDAC tissues. Specifically, spatial expression patterns and region-specific correlations for the *CXCL12–CXCR4*, *CCL2–CCR2* and *NT5E–CD8A* axes across all patients were examined. Spatial feature plots revealed a heterogeneous expression across tumour, stromal and immune-rich regions (Figure 4 and Supplementary Figure S2). *CXCL12, CXCR4* and *NT5E* expressions were predominantly enriched in tumour and stromal compartments, while *CD8A*, *CCR2* and *CCL2* showed broader distribution, with higher expression in immune-infiltrated areas. To quantify region-specific relationships, tissue sections were subdivided into spatial bins (500 × 500 pixels; ∼918 cells per bin), categorised as tumour-, stroma-, or immune--rich based on pathology-defined regions of interest (Supplementary Figure S3). Mean gene expression values within each bin were used to compute region-specific correlations.

**Figure 4.**
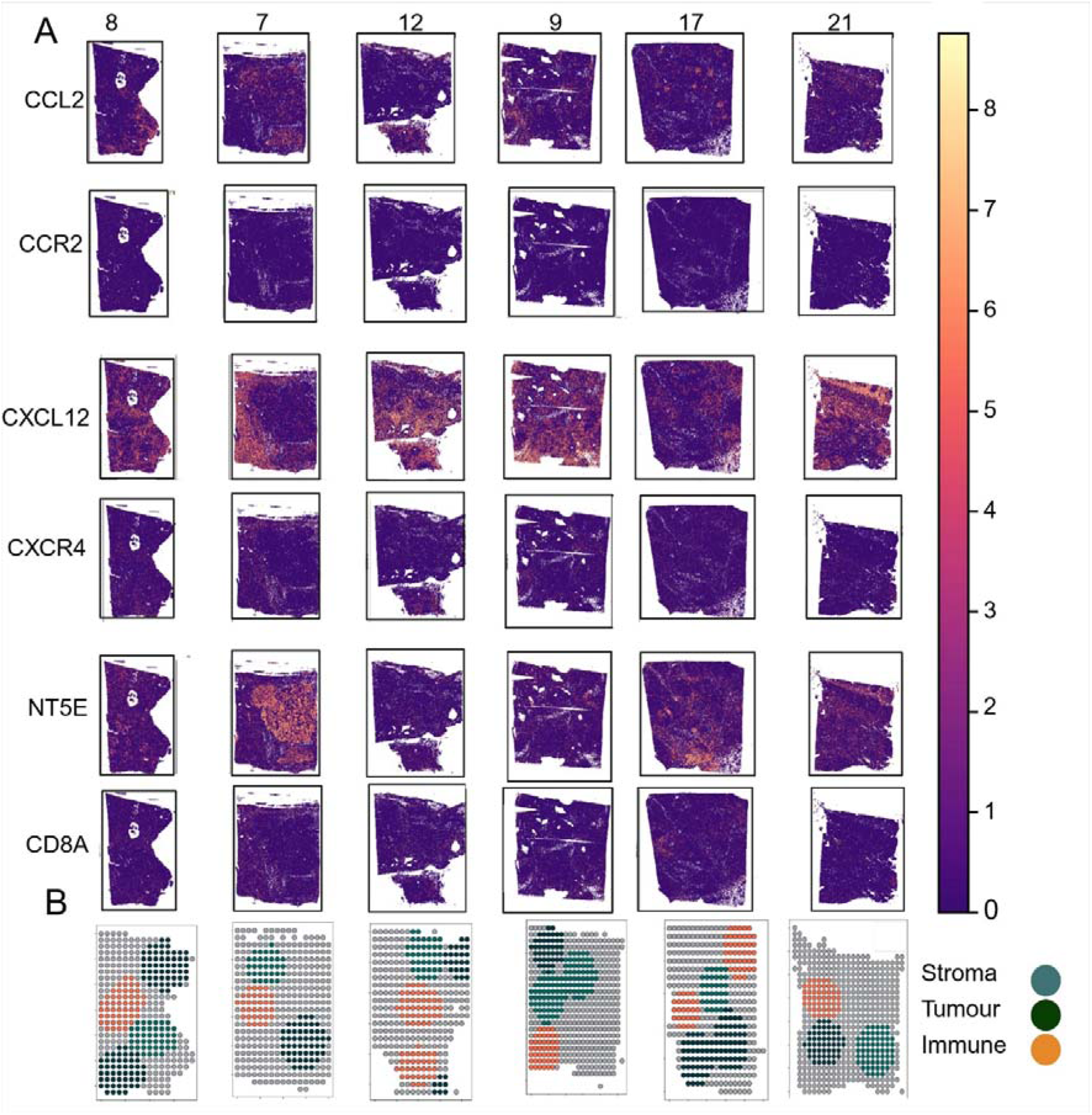
Spatial expression and compartmental annotation of chemokine signalling and immune-related genes in PDAC tissue sections. (A) Spatial feature plots showing log-transformed expression levels of *CCL2*, *CCR2*, *CXCL12*, *CXCR4*, *NT5E* and *CD8A* across all NAC and representative TN patient samples (NAC: 8, 7, 12 and 9, TN: 17 and 21). Warmer colours indicate higher expression. (B) Pathology-defined, color-coded regions of interest (ROIs) for each patient include tumour (dark green), stroma (light green) and immune-rich areas (orange). These annotated regions serve as a reference for interpreting the spatial distribution of the gene expression patterns shown in panel A.

*CXCL12* and *CXCR4* showed strong positive correlations across all regions in patients 8 and 7, with the strongest association observed in patient 8 (Figure 5A). In contrast, correlations were weak, absent, or reversed in patients 12 and 9, as well as in the immune region of TN patient 21. The *CCL2–CCR2* axis demonstrated positive correlations in tumour regions across all TN patients and NAC patients, with the strongest correlation observed in patient 8 (Figure 5B and Supplementary Figure S4C). Stromal correlations were negative or absent in NAC patients 8, 7 and 12, while patient 9 retained positive correlations across tumour and stromal compartments. NT5E and CD8A exhibited consistent negative correlations in tumour and stromal regions across patients 8 and 7, and TN samples (Figure 5C and Supplementary Figure S4B).

**Figure 5.**
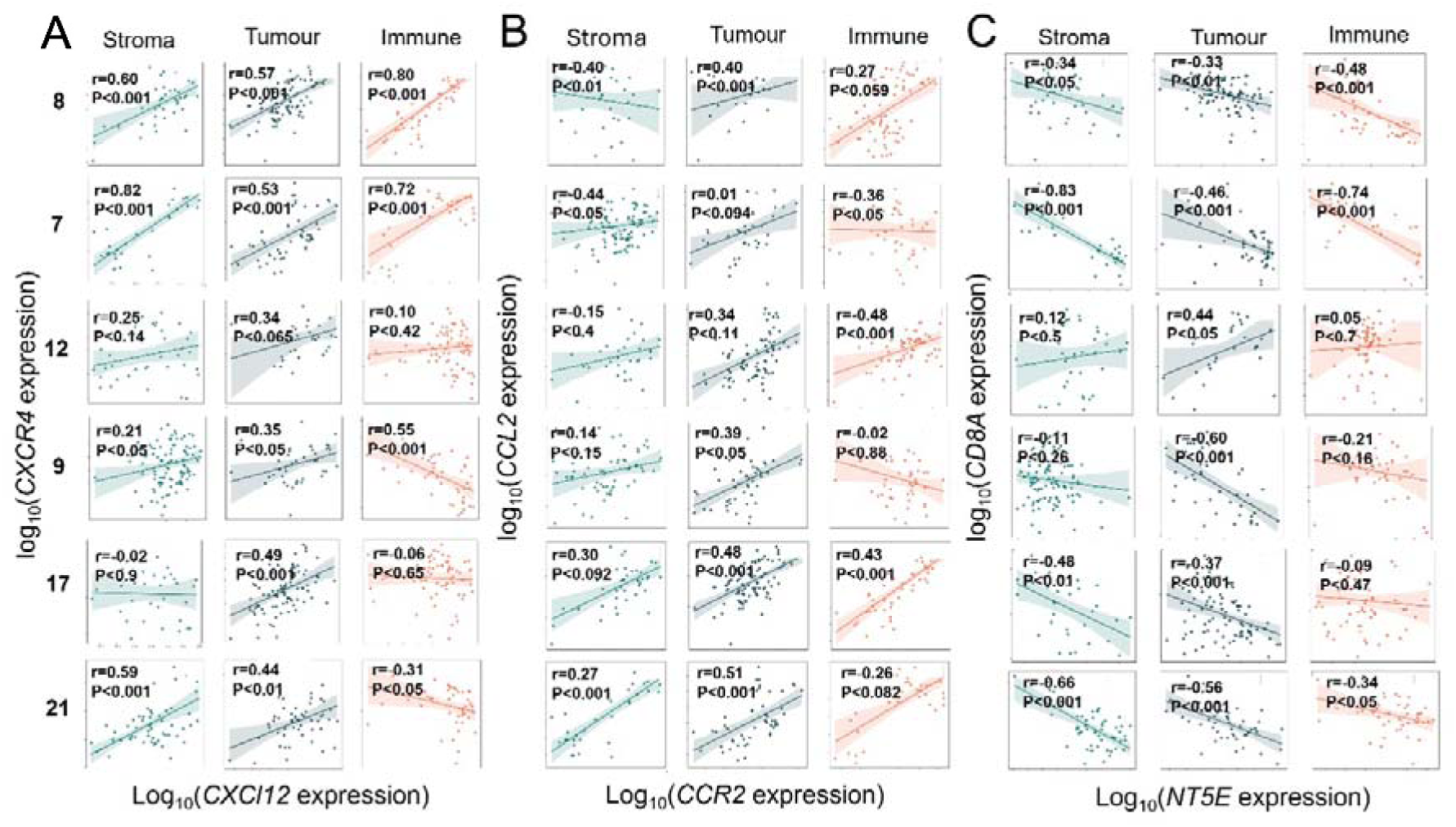
Region-specific spatial correlations of chemokine–receptor and immune suppression axes across PDAC patient tissues. (A) Correlation between *CXCL12* and *CXCR4* expression, (B) Correlation between *CCL2* and *CCR2* expression across regions, (C) Correlation between *NT5E* (CD73) and *CD8A* expression in stroma, tumour and immune-enriched regions for each patient. Each point represents a spatial bin (∼918 cells) and shaded areas indicate 95% confidence intervals. Pearson correlation coefficients (r) and corresponding p-values are shown. Correlations were calculated per patient and per region based on binned spatial expression values from annotated tissue compartments.

## Discussion

Neoadjuvant chemotherapy (NAC) is increasingly incorporated into PDAC management, yet its effects on immune–stromal architecture remains poorly defined. By integrating spatially annotated immune transcriptomics with *in situ* spatial profiling, we show that NAC induces substantial, regimen- and time-dependent immune–stromal reorganisation not apparent from bulk analyses. These data identify a transient, spatially defined window of immune remodelling, including dynamic modulation of *CXCL12–CXCR4* and *CCL2–CCR2* signalling axes, following chemotherapy.

Transcriptional segregation between treatment-naïve (TN) and NAC-treated tumours further supports widespread immune–stromal reprogramming. Clustering of some NAC samples with TN tumours, particularly from patients with shorter or undefined post-treatment intervals, suggests these alterations are temporally dynamic and partially reversible, consistent with previous single-cell and spatial transcriptomic studies (15–19). NAC was associated with coordinated upregulation of *CCL2, CXCL12, CD163, S100A12, CCL17* and *THBS1*, consistent with enhanced myeloid recruitment and immunosuppressive chemokine signalling (20–27). Increased expression of *CCL2* and *CXCL12*, implicating activation of the *CCR2* and *CXCR4* signalling axes, is consistent with their established roles in tumour progression and immune exclusion in PDAC (20, 28). However, emerging evidence suggests that chemokine function is context dependent, underscoring the importance of spatial localisation in interpreting these changes (29, 30).

Concurrent downregulation of *ITGB4*, *ITGA2*, and *NT5E* suggests partial disruption of tumour–stroma interactions and attenuation of fibrotic, immune-excluding programs (31–34). Increased *CD8A* expression further indicates that NAC may transiently enhance immune accessibility (6). However, persistent upregulation of *CXCL12* and *THBS1* suggests that T-cell functionality and retention may remain constrained. Together, these findings support a model in which chemotherapy partially remodels stromal barriers without fully overcoming immunosuppressive signalling (27, 28, 35).

Pathway enrichment analyses further support NAC-induced remodelling of the TME. Enrichment of mitotic and cell-cycle–related pathways may reflect the selective survival of chemotherapy-resistant tumour cell populations (36, 37). Although EMT pathways were enriched, the concurrent downregulation of key integrins involved in ECM anchoring suggests that NAC may disrupt EMT-associated tumour–stroma interactions rather than promote mesenchymal transition (38). Reactome and BioPlanet analyses similarly point to active ECM remodelling and engagement of stress-responsive transcriptional networks, consistent with an adaptive but potentially targetable post-chemotherapy state (39–42).

Spatial profiling demonstrated that immune accessibility was greatest in tumours resected shortly after NAC, whereas longer post-treatment intervals were associated with re-establishment of immune exclusion. These findings align with emerging evidence that cytotoxic therapies can induce immunogenic modulation of the tumour microenvironment, enhancing antigen presentation and T cell recruitment in a time-sensitive manner (19, 40). Elevated *CCL2* expression in non-malignant tissue has been reported in other cancers (41), and given the role of the *CCL2–CCR2* axis in immune cell trafficking (42), such gradients may redirect immune cells away from the tumour core, limiting effective anti-tumour immunity. Region-specific correlation analyses revealed regimen-dependent modulation of *CXCL12–CXCR4* and *CCL2–CCR2* coordination, supporting a temporally dynamic immune–stromal interaction landscape in the shorter resection intervals post NAC. Loss of coordination at longer intervals may reflect tissue recovery, receptor desensitisation or internalisation following sustained chemokine exposure (43, 44) or re-establishment of fibrosis-driven T-cell exclusion within the PDAC TME (45). Persistent negative correlation between NT5E and CD8A further supports a spatially restricted role for adenosine-mediated T-cell suppression (46).

These pathways are clinically relevant, as elevated *CXCL12–CXCR4*, *CCL2–CCR2*, and *NT5E* signalling have been consistently linked to poor outcomes in PDAC and other malignancies(20, 47–52). For instance, *CXCR4* has been shown to promote pancreatic cancer progression by regulating angiogenesis and lymphangiogenesis, as demonstrated through quantitative PCR–based analyses (53). Similarly, the *CCL2–CCR2* axis contributes to pathological angiogenesis, tumour cell survival and invasion, and the recruitment of immunosuppressive myeloid cells within the TME (54–56). In PDAC, extracellular ATP released by tumour cells is hydrolysed to adenosine through the *NT5E* (CD73) pathway, and this adenosine suppresses effector T-cell proliferation and cytotoxicity via the A_2A_ receptor, thereby impairing anti-tumour immunity (57, 58).

Our spatial analyses indicate that neoadjuvant chemotherapy reshapes the distribution and coordination of these pathways, supporting evaluation of immunomodulatory or stroma-directed strategies during defined post-chemotherapy intervals rather than after immune exclusion has re-emerged. This concept is supported by clinical evidence from the OPTIMIZE-1 trial, in which CD40 agonist–mediated stromal priming prior to chemotherapy enhanced immune activation and clinical outcomes (59–61). Together, these data suggest that stromal phenotype, immune architecture, and treatment timing collectively define therapeutic windows in PDAC. Future studies should evaluate neoadjuvant chemoimmunotherapy strategies with spatially resolved endpoints to optimise treatment sequencing and patient selection.

This study has several limitations. Its retrospective, non-randomised design introduces potential confounding by indication, and TN and NAC cohorts differed in treatment regimen, cycle number and timing of surgery, with some intervals unavailable, which may contribute to transcriptional heterogeneity. Gene expression analyses were limited to FFPE tissue and targeted panels, limiting coverage and interpretation to predefined pathways. Spatial transcriptomic analyses were performed on a small subset of representative samples and should be considered hypothesis-generating. Further, associations between chemokine expression and immune cell localisation are correlative, lacking protein-level validation, however using OnCorr, a specialised, open-source, web-based tool designed for pan-cancer mRNA–protein correlation analysis in pancreatic tissue, we observed strong concordance for *NT5E* (r = 0.85) and moderate concordance for *CD8A* (r = 0.63), supporting the significance of our findings. Finally, the study was not powered to link transcriptomic changes with clinical outcomes. Nevertheless, the reproducibility of spatial chemokine–receptor patterns across regions and patients underscores their biological relevance and provides a rationale for further mechanistic investigation.

In conclusion, NAC dynamically reshapes the PDAC immune microenvironment, with *CXCL12–CXCR4* and related pathways defining transient windows for potential therapeutic intervention. Timing of immunomodulatory and stroma-targeted strategies relative to chemotherapy may be critical to efficacy, warranting further mechanistic and clinical evaluation.

## Additional information

## Acknowledgments

This work was supported by the Cancer Institute NSW Career Development Fellowship (2020/CDF1093) and a Molecular Horizons Collaborative Grant, awarded to K.L. Vine-Perrow. We acknowledge the Garvan Genomic Facility and support from an Australian Government Research Training Program Scholarship and 2025 Millennium Science 10x Genomics Fellowship awarded to E. Minaei. The authors used GPT-4 to assist in improving the fluency of language during manuscript preparation; all content was subsequently reviewed and edited by the authors, who take full responsibility for the final version of the manuscript.

## Authors contribution

E.M. contributed to conceptualisation, methodology, investigation, analysis, visualisation, and writing (original draft, review and editing); D.C. and A.T. contributed to methodology, investigation, analysis, visualisation, and writing (review and editing); L.A.C. and K.H. contributed to investigation, data collection, and writing (review and editing); S.J.W., N.I., and D.W. contributed to methodology, investigation, and writing (review and editing); M.R., M.A. and R.S. contributed to supervision and writing (review and editing); EO contributed to writing (review and editing) and K.L.V. contributed to conceptualisation, supervision, resources, administration, investigation, and writing (review and editing).

## Ethics approval

This study was approved by the University of Wollongong and Illawarra Shoalhaven Local Health District Human Research Ethics Committee (UOW/ISLHD HREC 2020/ETH03297) and conducted in accordance with the Declaration of Helsinki and the National Statement on Ethical Conduct in Human Research (NHMRC, Australia).

## Data availability

The data generated in this study are available within the article and its supplementary data files. All statistical analyses were conducted in R (v4.2.0) and Python (v3.9) using standard packages and custom scripts available from Zenodo DOI: (10.5281/zenodo.17157651). All other raw data generated in this study are available from the corresponding author upon request.

## Competing interests

The authors declare no conflict of interest

## Footnotes

[1] Upper gastrointestinal (GI) cancer survival rates and statistics, National Cancer Institute, 2021 <u>National Trends in Cancer Death Rates Infographic - Annual Report to the Nation</u>

## Notes

https://zenodo.org/records/17157652

### Competing Interest Statement

The authors have declared no competing interest.

