## Supplementary material for "Neoadjuvant chemotherapy induces transient, regimen-specific immune–stromal reprogramming in pancreatic ductal adenocarcinoma": Supplementary materials.pdf

[1]School of Science, Faculty of Science Medicine and Health, University of Wollongong, Wollongong, NSW, Australia.
[2]Molecular Horizons, University of Wollongong, Wollongong, NSW, Australia.
[3]Illawarra Shoalhaven Local Health District, Wollongong, NSW, Australia.
[4]Illawarra Cancer Care Centre, Wollongong, NSW, Australia.
[5]Prince of Wales Hospital, School of Clinical Medicine, Randwick Clinical Campus, University of New South Wales, Sydney, NSW, Australia.
[6]New South Wales Health Pathology, Wollongong, NSW, Australia.
[7]New South Wales Health Pathology, Prince of Wales Hospital, Randwick, NSW, Australia.
[8]School of Clinical Medicine, UNSW Medicine & Health, University of New South Wales, Sydney, NSW, Australia.
[9] Department of Medicine, Memorial Sloan Kettering Cancer Center, New York, NY, USA
Weill Cornell Medicine, New York, NY, USA
** These co-authors contributed equally

#### Supplementary Method:

#### NanoString gene expression assays and data analysis

Up to 1 µg of purified RNA was run on the nCounter Sprint (NanoString Technologies, Seattle, USA) platform using the nCounter Human Pancancer Immune Profiling Assay (NanoString; 740 target genes, 30 housekeeping genes) as per the manufacturer's instructions. Quality control and analysis were performed using a Bioconductor package NanoStringNCTools [1], which flagged specimens with low binding density or other technical issues. These specimens were excluded from further data processing. Batch correction was conducted using the RUVg method from the RUVSeq Bioconductor package [2].

To ensure the integrity of the dataset, housekeeping genes exhibiting phenotype-specific associations were excluded using the Negative Binomial Generalised Linear Model (glm.nb) function within the RUVSeq pipeline [2]. The normalisation step involved testing all possible pairwise combinations with varying RUVg (remove unwanted variation using control genes) values. Normalised expression datasets were visualised using principal component analysis (PCA) and relative log expression (RLE) plots to identify outlier samples for further assessment [3]. After a rigorous QC and batch correction, a final cohort of specimens were proceeded to differential expression analysis, using DESeq2. Differentially expressed genes (DEGs) were identified based on $log_2$ fold changes and p-values adjusted for multiple testing using the Benjamini–Hochberg method to minimise false discovery rates [4]. The top DEGs were identified for comparisons between cohorts following the QC and normalisation steps.

#### Supplementary result and discussion:

#### Tissue-specific gene expression and pathway analysis revealed spatially divergent transcriptional programs

NanoString profiling of pathologist-selected micro-dissected *tumour* regions revealed a distinct transcriptional profile enriched for genes associated with cytotoxic and innate immune responses for NAC (Figure S1 and Supplementary Data file 1). Notably, *CD8A, CD3E, KLRB1, KLRC2* and *PRF1* were upregulated in NAC-treated tumour-rich areas (Figure S1B), which suggests increased infiltration or activation of $CD8^+$ T cells and natural killer cells, consistent with prior reports linking cytotoxic chemotherapy to immunogenic cell death and enhanced T cell priming in PDAC [5]. In parallel, the study also observed increased expression of *CCL5* and *CXCL12*, chemokines that regulate T cell and myeloid cell trafficking

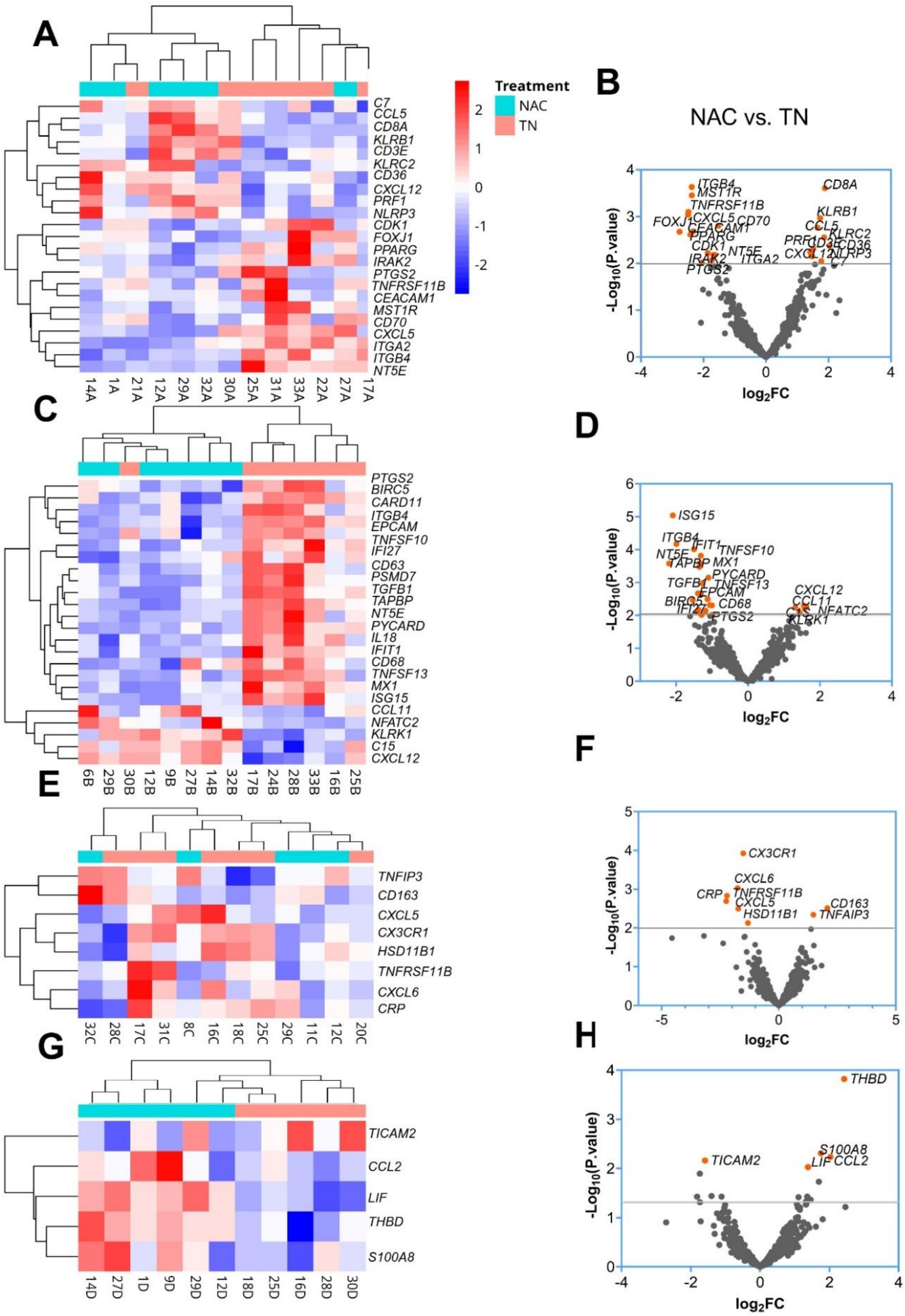


**Figure S1. Compartment-specific Heatmap and Volcano plots in PDAC following neoadjuvant chemotherapy.** Heatmap showing unsupervised clustering and volcano plot of top differentially expressed immune-related genes in (A–B) tumour-rich regions (C–D) stroma-rich compartments, (E–F) immune-rich regions and (G–H) tumour-adjacent normal tissue comparing neoadjuvant-treated (NAC, cyan) and treatment-naïve (TN, pink) samples (p-value< 0.01, Log2 Foldchange > 1 and < -1).

[6]. CCL5/CCR5 signalling is widely associated with tumour progression, driving immunosuppression through the recruitment of regulatory T cells, myeloid-derived suppressor cells and tumour-associated macrophages [7, 8]. However, in KPC models, CD40 agonist antibody induces CCL5 production by intra-tumoural myeloid cells, where it instead facilitates $CD4^+$ T cell infiltration and supports anti-tumour immunity [9]. On the other hand, the upregulation of *CXCL12* in NAC, consistent with the bulk tumour analysis with its role in T cell exclusion and immunotherapy resistance [10], reinforce the spatial complexity of chemokine function in the TME. On the other hand, NAC-treated tumours showed a reduced expression of *ITGB4*, *ITGA2*, *CEACAM1* and *MST1R* genes in tumour-rich areas (Figure S1B), linked to cell adhesion, stromal remodelling and immunosuppressive signalling [11-15]. Pathway enrichment analysis using MSigDB Hallmark databases revealed that EMT was among the most differentially enriched pathways in tumour-rich tissue following neoadjuvant chemotherapy, suggesting enhanced tumour cell plasticity and stress adaptation (Table S3). EMT enrichment was supported by concurrent activation of KEGG pathways involved in ECM–receptor interaction and regulation of the actin cytoskeleton, highlighting a coordinated transcriptional program affecting cell–matrix adhesion and cytoskeletal remodelling (Table S4).

NanoString gene expression analysis within the *stromal* compartment revealed that NAC-treated stromal regions displayed a markedly different transcriptional profile compared to TN tissues (Figure S1C and Supplementary Data file 1). NAC profoundly reprogrammed stromal regions, with broad downregulation of inflammatory, interferon and fibrotic gene programs. Key mediators of desmoplasia and immune exclusion, including *PTGS2, TGFB1, NT5E* (CD73), *ITGB4*, *TNFSF10* and interferon-stimulated genes (*ISG15, MX1, IFI27, TAPBP*), were suppressed in NAC-treated stromal-rich compartments (Figure S1D), suggesting reduced stromal immunogenicity and structural barriers [16-21]. *NT5E* encodes CD73 (5′-

nucleotidase), a key regulator of extracellular adenosine production that contributes to T cell suppression and immune evasion in PDAC [22]. This structural and metabolic reprogramming may, in part, explain the significant increase in *CD8A* and *CD3E* gene expression, markers of cytotoxic T cells, observed in NAC tumours compared to TN tumours. Select chemokines and immune modulators such as *CXCL12, CCL11, NFATC2, C1S* and *KLRK1* were upregulated in stromal regions of NAC patients, potentially maintaining immune exclusion and regulatory phenotypes [23-25]. Notably, the reduced expression of structural immune-exclusion markers (*NT5E, ITGB4*), concurrent with upregulation of exclusion-associated chemokines such as *CXCL12*, suggests a dual-phase stromal reprogramming: one that permits immune cell infiltration while maintaining local immunosuppression through soluble mediators. Pathway enrichment analysis confirmed the suppression of interferon alpha and gamma responses, with downstream effects on antiviral and immune-priming pathways, underscoring the spatial heterogeneity of stromal responses post-NAC (Tables S5, S6).

In immune rich regions, bulk profiling revealed that NAC induced a paradoxical immune signature (Figure S1E and Supplementary datafile). Inflammatory and chemotactic transcripts such as *CXCL5, CXCL6* and *CX3CR1* were downregulated, while immunosuppressive markers *CD163* (M2-like macrophages) and *TNFAIP3* (negative regulator of NF-κB) were upregulated. (Figure 2F). Pathway enrichment analyses identified interleukin-1 regulation of extracellular matrix, Oncostatin M signalling and chemokine–receptor interactions as key altered pathways, indicating rewiring of cytokine–matrix networks that control immune cell positioning [26-28] (Table S7). Additional enrichment of glucocorticoid biosynthesis, angiogenesis, tumour necrosis factor (TNF)-α signalling and hypoxia-related programs supports a shift toward immunosuppression and stress adaptation [29-31]. EMT and xenobiotic metabolism pathways were also enriched, suggesting therapy-induced stromal plasticity [32-34] (Table S8). Together, these findings reveal that NAC remodels immune-rich regions by dampening inflammatory responses while simultaneously reinforcing immunoregulatory and matrix-associated signalling networks.

Finally, in tumour-adjacent normal tissue, even histologically normal regions bordering the tumour exhibited marked transcriptional reprogramming post-NAC (Figure S1G and Supplementary datafile). Genes involved in immune activation and tissue stress, including *THBD, S100A8, CCL2* and *LIF*, were significantly upregulated, implicating vascular modulation, myeloid recruitment and stress-adaptive signalling [35-38] (Figure S1H). Pathway enrichment analyses identified the unfolded protein response (UPR), PERK-regulated gene expression, coagulation and interleukin-1 and MSP/RON receptor signalling, indicating endoplasmic reticulum stress, early immune surveillance and matrix adaptation. These peripheral changes suggest that normal tissue may engage low-level regulatory programs in response to chemotherapy, potentially influencing tumour recurrence or systemic immune outcomes [39-42] (Tables S9 and S10). Although none of these pathways were significantly enriched after multiple testing correction, their presence suggests that normal tissues may not be fully quiescent but instead engage low-level regulatory programs related to stress adaptation, cytokine signalling and matrix homeostasis. These findings raise the possibility that even tumour-adjacent normal tissues may exhibit early immunoregulatory reprogramming that could influence tumour progression or response to therapy.

Collectively, these findings demonstrate that NAC induces compartment-specific immune and structural reprogramming across tumour, stromal, immune-rich and adjacent normal tissues. The progressive nature of these changes from the tumour core to the periphery highlights the interconnectedness of the PDAC microenvironment and underscores the need for integrative single cell spatial analyses to disentangle permissive versus suppressive immune dynamics and to guide rational combinations of NAC with targeted immunomodulatory therapies.

### Supplementary Tables and Figures

##### Table S1 Pooled Tissue Reactome_Pathways_2024

| NAME | P-value | Adj p-value | Odds ratio | Combined score |
|---|---|---|---|---|
| **Non-integrin membrane-ECM interactions** | 0.0026 | 0.1826 | 15.89 | 94.51 |
| **Syndecan interactions** | 0.0026 | 0.1826 | 15.89 | 94.51 |
| **Mitotic prometaphase** | 0.0033 | 0.1826 | 61.22 | 349.58 |
| **Resolution of sister chromatid cohesion** | 0.0033 | 0.1826 | 61.22 | 349.58 |
| **Nicotinate metabolism** | 0.0064 | 0.1826 | 30.57 | 154.01 |
| **Metabolism of water-soluble vitamins and cofactors** | 0.0064 | 0.1826 | 30.57 | 154.01 |
| **Laminin interactions** | 0.0105 | 0.1826 | 20.35 | 92.56 |
| **Mitotic anaphase** | 0.0155 | 0.1826 | 15.24 | 63.47 |
| **Mitotic metaphase and anaphase** | 0.0155 | 0.1826 | 15.24 | 63.47 |

##### Table S2 Pooled Tissue BioPlanet_2019

| NAME | P-value | Adj p-value | Odds ratio | Combined score |
|---|---|---|---|---|
| **BDNF signalling pathway** | 9.85E-5 | 0.0279 | 11.40 | 10.518 |
| **FRA pathway** | 0.0048 | 0.2751 | 11.88 | 63.21 |
| **AP-1 transcription factor network** | 0.0061 | 0.2751 | 6.88 | 35.05 |
| **Regulation of NFAT transcription factors** | 0.0083 | 0.2751 | 6.20 | 29.70 |
| **TSH regulation of gene expression** | 0.0100 | 0.2751 | 8.60 | 39.55 |
| **MSP/RON receptor signalling pathway** | 0.0105 | 0.2751 | 20.35 | 92.56 |
| **Axon guidance** | 0.0125 | 0.2751 | 5.40 | 23.65 |
| **ECM-receptor interaction** | 0.0148 | 0.2751 | 7.26 | 30.58 |
| **FOXM1 transcription factor network** | 0.0212 | 0.2751 | 12.17 | 46.86 |
| **Pertussis toxin-insensitive CCR5 signalling in macrophage** | 0.0278 | 0.2751 | 10.13 | 36.29 |

##### Table S3 Tumour Tissue MSigDB_2020

| NAME | P-value | Adj p-value | Odds ratio | Combined score |
|---|---|---|---|---|
| **Epithelial mesenchymal transition** | 0.0119 | 0.2690 | 5.51 | 24.43 |
| **Adipogenesis** | 0.0299 | 0.2691 | 9.52 | 33.43 |
| **Glycolysis** | 0.0299 | 0.2691 | 9.52 | 33.43 |
| **Apical junction** | 0.0487 | 0.3240 | 4.27 | 12.90 |
| **Myogenesis** | 0.0600 | 0.3240 | 6.03 | 16.95 |
| **Fatty acid metabolism** | 0.0917 | 0.4126 | 16.02 | 38.28 |
| **Mitotic spindle** | 0.1204 | 0.4645 | 10.67 | 22.58 |
| **Spermatogenesis** | 0.1753 | 0.5720 | 6.38 | 11.11 |
| **E2f targets** | 0.2015 | 0.5720 | 4.55 | 6.740 |
| **Cholesterol homeostasis** | 0.2269 | 0.5720 | 4.55 | 6.740 |

**Table S4 KEGG 2021 Human**

| NAME | P-value | Adj p-value | Odds ratio | Combined score |
|---|---|---|---|---|
| **PPAR signaling pathway** | 0.002798 | 0.2881 | 67.24 | 395.29 |
| **ECM-receptor interaction** | 0.009722 | 0.5007 | 8.69 | 40.25 |
| **AMPPK signaling pathway** | 0.02371 | 0.6296 | 11.13 | 41.64 |
| **Arrhythmogenic right ventricular cardiomyopathy** | 0.03666 | 0.6296 | 8.32 | 27.51 |
| **Regulation of actin cytoskeleton** | 0.04424 | 0.6296 | 4.46 | 13.91 |
| **Dilated cardiomyopathy** | 0.05175 | 0.6296 | 6.64 | 19.66 |
| **Hypertrophic cardiomyopathy** | 0.06 | 0.6296 | 6.03 | 16.95 |
| **Fat digestion and absorption** | 0.06206 | 0.6296 | 32.09 | 89.2 |
| **Ovarian steroidogenesis** | 0.06206 | 0.6296 | 32.09 | 89.2 |

**Table S5 Stromal Tissue MSigDB_2020**

| NAME | P-value | Adj p-value | Odds ratio | Combined score |
|---|---|---|---|---|
| **Interferon gamma response** | 5.37E-04 | 0.0145 | 5.283333 | 39.78041 |
| **Interferon alpha response** | 0.021491 | 0.290128 | 4.506667 | 17.30615 |
| **G2-M checkpoint** | 0.0397 | 0.357296 | 7.931818 | 25.59134 |
| **Myogenesis** | 0.064796 | 0.376222 | 5.743802 | 15.71802 |
| **Epithelial mesenchymal transition** | 0.070721 | 0.376222 | 3.592593 | 9.516833 |
| **MYC targets V1** | 0.095549 | 0.376222 | 15.30435 | 35.93632 |
| **Mitotic spindle** | 0.125407 | 0.376222 | 10.18841 | 21.15304 |
| **Protein secretion** | 0.125407 | 0.376222 | 10.18841 | 21.15304 |
| **Unfolded protein response** | 0.125407 | 0.376222 | 10.18841 | 21.15304 |

**Table S6 Stromal Tissue BioPlanet_2019**

| NAME | P-value | Adj p-value | Odds ratio | Combined score |
|---|---|---|---|---|
| **Antiviral mechanism by interferon-stimulated genes** | 0.004342 | 0.520599 | 12.46 | 67.79 |
| **Proteins and DNA sequences in cardiac structures** | 0.019681 | 0.520599 | 12.7 | 50.06 |
| **Interferon alpha/beta signalling** | 0.028622 | 0.520599 | 4.07 | 14.49 |
| **Interleukin-11 pathway** | 0.032405 | 0.520599 | 9.07 | 31.13 |
| **DNA replication** | 0.032405 | 0.520599 | 9.07 | 31.13 |
| **Phase I of biological oxidations: functionalization of compounds** | 0.032877 | 0.520599 | Infinity | Infinity |
| **Purine catabolism** | 0.032877 | 0.520599 | Infinity | Infinity |
| **Pyrimidine catabolism** | 0.032877 | 0.520599 | Infinity | Infinity |
| **Aurora b signalling** | 0.032877 | 0.520599 | Infinity | Infinity |

**Table S7 Immune Tissue BioPlanet_2019**

| NAME | P-value | Adj p-value | Odds ratio | Combined score |
|---|---|---|---|---|
| **Interleukin-1 regulation of extracellular matrix** | 3.90E-05 | 0.0022 | 27.68293 | 281.0541 |
| **Oncostatin M** | 0.0019 | 0.0564 | 12.37037 | 76.99442 |
| **FSH regulation of apoptosis** | 0.0030 | 0.0581 | 15.44444 | 89.42324 |
| **Binding of chemokines to chemokine receptors** | 0.0105 | 0.0780 | 9.474419 | 43.16643 |
| **Steroid hormone biosynthesis** | 0.0109 | 0.0780 | Infinity | Infinity |
| **Steroid hormone, vitamin a, and vitamin d metabolism** | 0.0109 | 0.0780 | Infinity | Infinity |
| **Glucocorticoid and mineral corticoid metabolism** | 0.0109 | 0.0780 | Infinity | Infinity |
| **Glucocorticoid biosynthesis** | 0.0109 | 0.0780 | Infinity | Infinity |
| **Peptide G-protein coupled receptors** | 0.0181 | 0.1151 | 7.573585 | 30.35119 |

**Table S8 Immune Tissue MSigDB_2020**

| NAME | P-value | Adj p-value | Odds ratio | Combined score |
|---|---|---|---|---|
| **Epithelial mesenchymal transition** | 0.0030 | 0.0305 | 15.444 | 89.423 |
| **Xenobiotic metabolism** | 0.0116 | 0.0584 | 16.857 | 75.000 |
| **KRAS signalling up** | 0.0820 | 0.2119 | 5.2635 | 13.159 |
| **Angiogenesis** | 0.0847 | 0.2119 | 14.591 | 36.009 |
| **TNF-alpha signalling via NF-KB** | 0.1311 | 0.2402 | 3.8888 | 7.8992 |
| **Hypoxia** | 0.1441 | 0.2402 | 7.7912 | 15.090 |
| **Apical junction** | 0.2613 | 0.3734 | 3.8241 | 5.1310 |
| **Complement** | 0.4673 | 0.5841 | 1.7671 | 1.3444 |
| **Interferon gamma response** | 0.6115 | 0.6304 | 1.1464 | 0.5636 |
| **Inflammatory response** | 0.6304 | 0.6304 | 1.0850 | 0.5005 |

**Table S9 Tumour adjacent Normal Tissue BioPlanet_2019**

| NAME | P-value | Adj p-value | Odds ratio | Combined score |
|---|---|---|---|---|
| **FRA pathway** | 2.51E-05 | 0.0014 | 134.43 | 1423.8 |
| **Unfolded protein response** | 0.0068 | 0.1347 | Infinity | Infinity |
| **PERK-regulated gene expression** | 0.0068 | 0.1347 | Infinity | Infinity |
| **Coagulation common pathway** | 0.0136 | 0.2009 | 181 | 777.07 |
| **Low-density lipoprotein (LDL) pathway during atherogenesis** | 0.0204 | 0.2009 | 90.37 | 351.6036 |
| **Diabetes pathways** | 0.0204 | 0.2009 | 90.37 | 351.60 |
| **Interleukin-1 signalling pathway** | 0.0329 | 0.2255 | 10.57 | 36.06 |
| **MSP/RON receptor signalling pathway** | 0.0338 | 0.2255 | 45.06 | 152.54 |
| **Interleukin-1 regulation of extracellular matrix** | 0.0344 | 0.2255 | 10.318 | 34.76 |

**Table S10 Tumour adjacent Normal Tissue MSigDB_2020**

| NAME | P-value | Adj p-value | Odds ratio | Combined score |
|---|---|---|---|---|
| **Unfolded protein response** | 0.0271 | 0.1826 | 60.166 | 216.93 |
| **Angiogenesis** | 0.0537 | 0.1826 | 25.642 | 74.964 |
| **TNF-alpha signalling via NF-KB** | 0.0548 | 0.1826 | 7.8128 | 22.688 |
| **Inflammatory response** | 0.1060 | 0.2652 | 5.1566 | 11.568 |
| **P53 pathway** | 0.1422 | 0.2844 | 8.3809 | 16.344 |
| **Allograft rejection** | 0.183 | 0.3059 | 3.5362 | 5.9950 |
| **Coagulation** | 0.2183 | 0.3119 | 5.0808 | 7.7312 |
| **KRAS signalling up** | 0.2731 | 0.3414 | 3.8693 | 5.0213 |
| **IL-2/STAT5 signalling** | 0.3298 | 0.3664 | 3.0454 | 3.3780 |
| **Interferon gamma response** | 0.4455 | 0.4455 | 2.0156 | 1.6295 |

**Table S11 Clinical characteristics of PDAC patients included in Xenium spatial transcriptomics analysis**

| Patient | Treatment | Type of treatment | Gender | Interval between the last cycle of NAC and surgery (weeks) | Number of cycles |
|---|---|---|---|---|---|
| **7** | NAC | FOLFIRINOX | M | Not known | 6 |
| **8** | NAC | FOLFIRINOX | F | 3 | 7 |
| **9** | NAC | Gem/nab-paclitaxel | M | 12 | 12 |
| **12** | NAC | FOLFIRINOX | F | 12 | 6 |
| **17** | TN | nil | F | NA | NA |
| **21** | TN | nil | F | NA | NA |
| **24** | TN | nil | M | NA | NA |
| **25** | TN | nil | M | NA | NA |

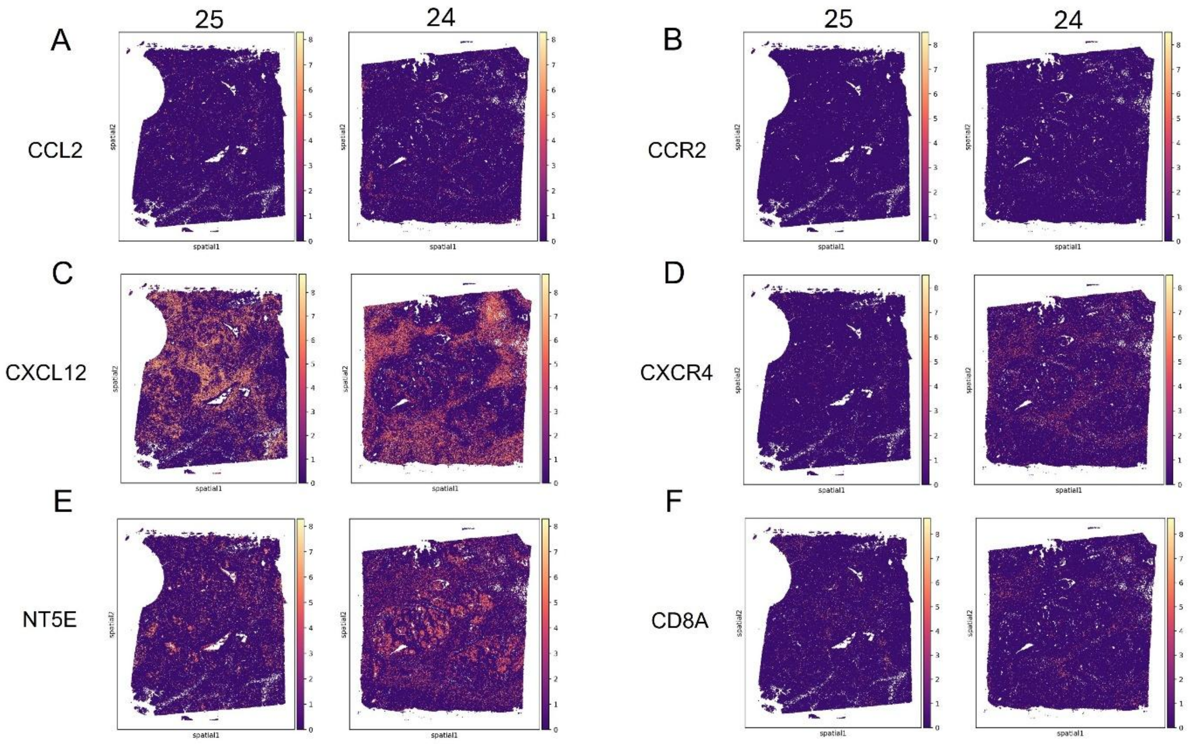


**Figure S2. Spatial expression of chemokines, receptors, and immune markers in treatment-naïve PDAC patients (patients 24 and 25).** Spatial transcriptomic feature plots show log-transformed expression levels of (A) *CCL2*, (B) *CCR2*, (C) *CXCL12*, (D) *CXCR4*, (E) *NT5E*, and (F) *CD8A* across annotated tumour tissue sections from two treatment-naïve PDAC patients. High expression is indicated by yellow/white and low expression by purple (log-normalized scale).

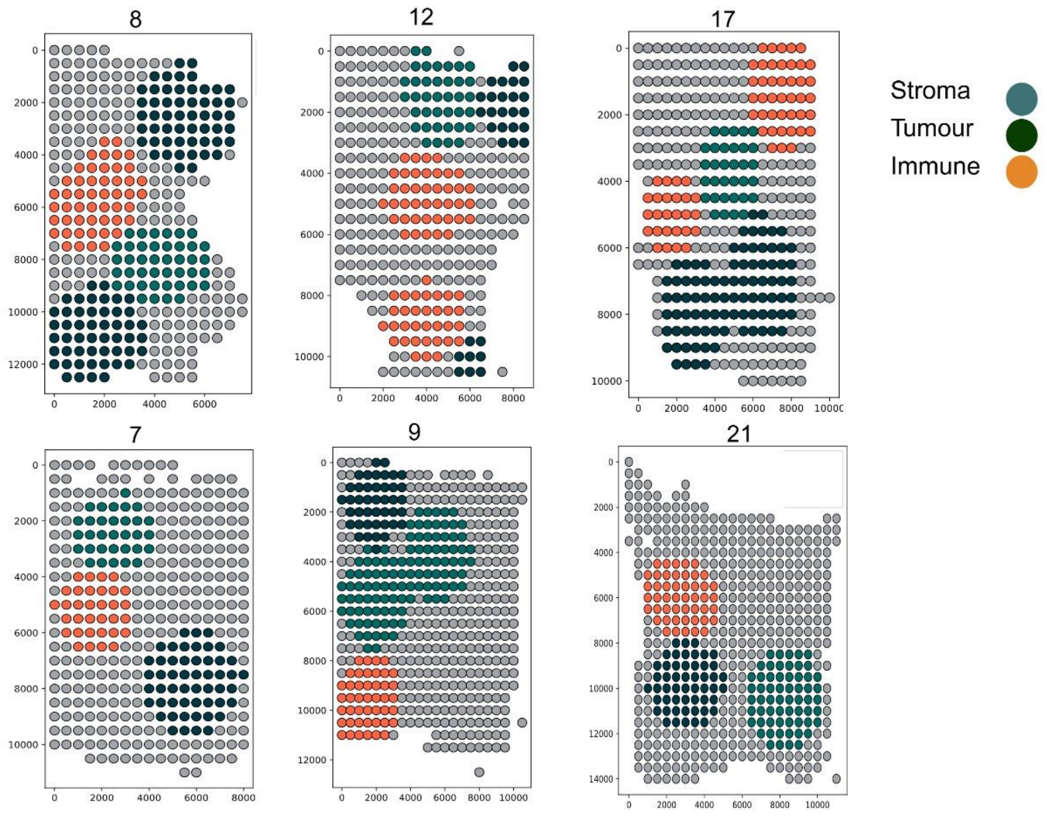


**Figure S3.** Spatial binning of each tissue section into 500 × 500 pixel regions (~918 cells per bin), color-coded based on the dominant cell type: tumours (dark green), stroma (light green), or immune-rich (orange). These annotated bins were used for downstream region-specific correlation analysis in Figures 6A–C and Supplementary Figure S3.

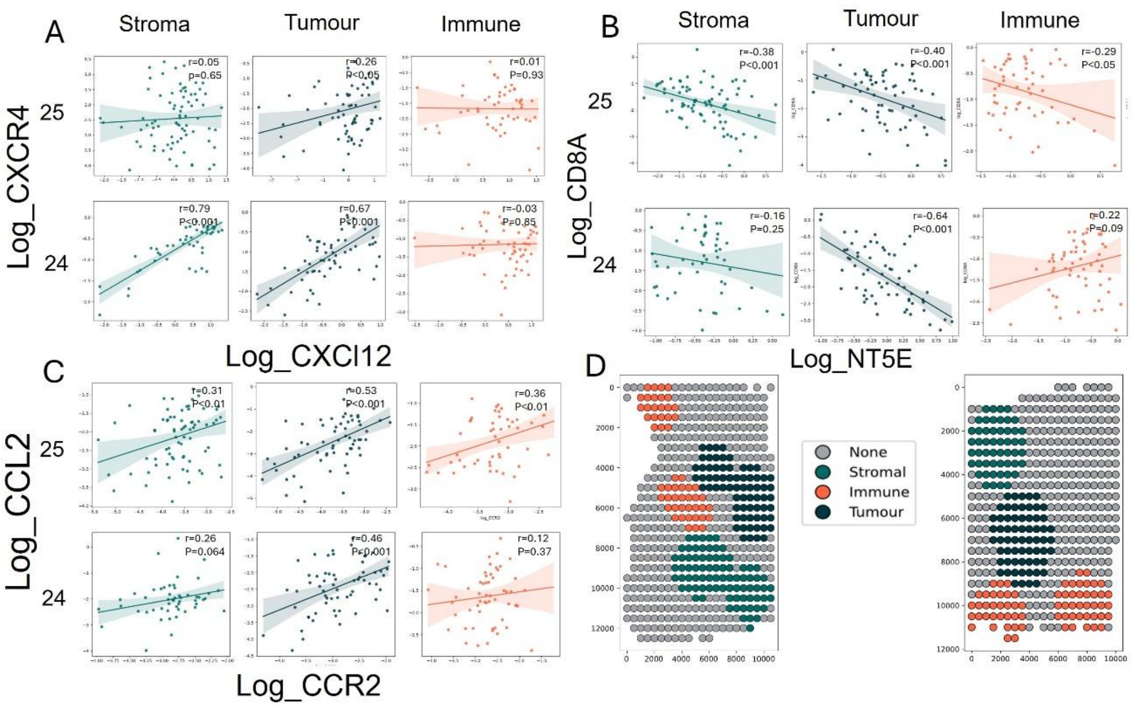


**Figure S4. Region-specific spatial correlations of chemokine-receptor and immunomodulatory axes in treatment-naïve PDAC patients (24 and 25).** (A–C) Scatterplots showing region-specific spatial correlations in stromal, tumour, and immune-enriched regions for:(A) *CXCL12–CXCR4* axis,(B) *NT5E–CD8A* axis,(C) *CCL2–CCR2* axis. Each point represents a spatial bin (~918 cells per bin), colored by region. Correlation coefficients (r) and p-values are shown for each region and patient. (D) Spatial bin annotation maps for patients 24 and 25, showing categorization of each bin into stromal (green), immune (orange), and tumour (blue) based on dominant cell type composition.
